# Safeguarding Privacy in Genome Research: A Comprehensive Framework for Authors

**DOI:** 10.1101/2024.09.20.614092

**Authors:** Maryam Ghasemian, Lynette Hammond Gerido, Erman Ayday

**Affiliations:** Case Western Reserve University, Cleveland, OH

**Author notes:** Corresponding Authors: Lynette Hammond Gerido, Erman Ayday.

## Abstract

As genomic research continues to advance, sharing of genomic data and research outcomes has become increasingly important for fostering collaboration and accelerating scientific discovery. However, such data sharing must be balanced with the need to protect the privacy of individuals whose genetic information is being utilized. This paper presents a bidirectional framework for evaluating privacy risks associated with data shared (both in terms of summary statistics and research datasets) in genomic research papers, particularly focusing on re-identification risks such as membership inference attacks (MIA). The framework consists of a structured workflow that begins with a questionnaire designed to capture researchers’ (authors’) self-reported data sharing practices and privacy protection measures. Responses are used to calculate the risk of re-identification for their study (paper) when compared with the National Institutes of Health (NIH) genomic data sharing policy. Any gaps in compliance help us to identify potential vulnerabilities and encourage the researchers to enhance their privacy measures before submitting their research for publication. The paper also demonstrates the application of this framework, using published genomic research as case study scenarios to emphasize the importance of implementing bidirectional frameworks to support trustworthy open science and genomic data sharing practices.

## Introduction

Large collections of human genomes representing centuries of data are transforming research and clinical care, opening new frontiers in our understanding of human health and disease. Large-scale studies, such as Genome-Wide Association Studies (GWAS) [1], have provided invaluable insights by linking genetic variations to specific traits and conditions. As the volume of genomic data continues to grow, so does the need for sharing this data among researchers to facilitate further discoveries and replicate findings.

However, this new era of genomic research and medical informatics brings significant challenges for clinical informaticians and quantitative health researchers. They must adapt their privacy-preserving practices and policies in line with the rapidly evolving landscape of information technologies. While many researchers are aware of technological advancements and the benefits of data sharing, few consider the increased privacy risks driven by academic pressures to publish. Without additional safeguards and risk-informed practices in publication and data sharing, research outputs may fail to effectively protect individual privacy when disseminating findings and sharing genomic data.

Given the sensitive nature of genomic data, there are substantial privacy challenges, particularly the risk of reidentifying individuals from shared datasets. Addressing these concerns requires robust frameworks that balance the imperative for data sharing with the need to maintain confidentiality and trust.

To address these concerns, we propose a bidirectional framework for evaluating privacy risks associated with data shared (both in terms of summary statistics and research datasets) in genomic research papers. Because privacy protections must be considered from a case-by-case approach, our framework requires input from the researchers (i.e., authors of such papers). This bidirectional framework integrates a questionnaire to assess researchers’ data sharing practices with a structured workflow that compares these practices against NIH guidelines [2]. Additionally, the framework includes a Membership Inference Attack (MIA) analysis to quantify the re-identification risks associated with the shared data.

### Significance

Publishing results of genomic research and sharing genomic data enhances the reproducibility of scientific findings through rigor and transparency. However, researchers and publishers must understand the varying contexts in which the content may be used. It is also essential to balance the priorities of research while maintaining the trust, particularly in a field as sensitive as genomic research. Inadequate privacy protections during peer review and publication can lead to disparities and diminish public confidence, especially when third parties can potentially trace genomic data back to individuals.

A particularly concerning type of privacy risk is membership inference. Membership Inference Attacks (MIAs) are conducted by malicious actors to determine if a specific individual’s data was included in a dataset used to train a model or to re-identify encrypted or anonymized information through data mining techniques. Such re-identification can have severe ethical, legal, and social implications that may affect not only the individual but also their relatives, with potential consequences cascading throughout their kinship network.

### Objectives

To mitigate the risks mentioned above, we present a bidirectional framework for evaluating privacy risks in genomic research papers considering how the authors share the outcome of their research (both in terms of summary statistics and research datasets) in such papers. This framework is applied to publicly available genomic research articles as case studies. These case studies highlight the critical need for researchers to implement and continuously enhance privacy protection measures to mitigate risks and uphold the ethical standards of genomic research. The findings underscore the importance of balancing data accessibility with the protection of individual privacy, ensuring that the benefits of genomic research are realized without compromising the confidentiality of research participants.

### Overview of the Bidirectional Framework

Our Bidirectional Framework is summarized in the following three steps, which will be discussed later in the ‘Methods’ section.

1. Questionnaire: Researchers (i.e., authors) share the privacy characteristics of the study.
2. Comparative Analysis: An algorithm compares the study characteristics against NIH Genomic Data Sharing Policy.
3. Feedback: Summary of Analysis & Suggested Actions Returned to the Researchers.

### Background

In this section we review some background related to genomic research as well as privacy threads.

### Genomic Background

The human genome consists of a sequence of nucleotides—A, T, C, and G—forming approximately 3 billion nucleotide pairs. Among these, 99.9% of the genome is identical between any two individuals, with the remaining 0.1% accounting for genetic variations. The most common type of genetic variation is single nucleotide polymorphisms (SNPs), which result from differences in a single nucleotide and are typically biallelic, meaning they have two possible forms or alleles at a specific genetic location.

Most genomics-related studies such as Genome-Wide Association Studies (GWAS) are a prevalent method used to identify genetic variations associated with particular traits or phenotypes. In a typical GWAS approach, known as a case-control study, researchers compare the genomes of individuals with a specific trait (cases) to those without it (controls). The study aims to identify SNPs that correlate with the trait of interest. In this context, SNPs are treated as biallelic markers with values 0, 1, or 2, indicating the number of minor alleles present. The data from these studies is often summarized in contingency tables that display the distribution of SNP values across cases and controls.

However, sharing data and statistical results from genomic studies, such as p-values, odds ratios, and minor allele frequencies (MAFs) for significant SNPs poses privacy risks. Despite data anonymization efforts, it is possible to infer individual participation or re-identify individuals within a dataset, particularly when sharing detailed SNP statistics. Even with data anonymization efforts, there remains a risk of inferring individual participation or re-identifying individuals within a dataset, particularly when detailed statistical information is shared. Some studies choose to share only summary statistics, while others share both the underlying data and associated statistics. These varying practices highlight the need for robust frameworks to assess and mitigate re-identification risks. This paper focuses on developing a systematic approach to evaluate privacy risks associated with the publication that practices some type of genomics-related data and/or statistics sharing.

### Privacy Threats

In the context of genomic research, several types of privacy attacks can exploit research findings and partially shared datasets. The most prominent among these are membership inference attacks [3, 4], attribute inference attacks [5, 6, 7, 8], and deanonymization attacks [9, 10, 11]. Each of these attacks poses distinct privacy risks to individuals whose data is included in genomic studies.

Membership inference attacks involve an adversary attempting to determine whether a specific individual’s genomic data is part of a research dataset. This attack is particularly concerning when datasets are shared publicly or even within controlled environments, as the attacker uses statistical clues or patterns in the dataset to infer membership. For example, if an attacker knows some characteristics of a target individual’s genome, they can compare this information against the dataset to determine whether that individual was included in the study. Previous studies have shown that aggregate statistics published as part of GWAS are especially vulnerable to membership inference attacks [3, 4, 12, 13].Successful membership inference can have serious privacy implications, especially in cases where the presence of an individual in a dataset could reveal sensitive information about their health or genetic traits.

Attribute inference attacks go a step further by aiming to deduce additional private information about an individual based on the data already revealed. In these attacks, the adversary uses the observed genetic data to infer unknown attributes, such as predispositions to certain diseases or other personal traits. For instance, if part of an individual’s genome is known, an attacker could use this information to predict other genomic attributes that are not explicitly shared. This type of attack becomes particularly relevant when genomic data is linked to an individual or a specific group, allowing attackers to uncover details that the data subjects might not have consented to share.

Deanonymization attacks are another critical threat in the domain of genomic data sharing. The primary goal of these attacks is to re-identify individuals from anonymized datasets. Even when genomic data is stripped of direct identifiers, such as names or addresses, it can often still be linked back to an individual using auxiliary information. This auxiliary information could include observable phenotypic traits (e.g., eye color or height) or known segments of an individual’s genome. Deanonymization is especially dangerous because once an individual’s identity is linked to their genomic data, it may lead to unintended consequences, including discrimination, stigmatization, or misuse of personal health information.

While attribute inference attacks are contingent upon the ability to link genomic data to specific individuals, membership inference and deanonymization attacks are of particular concern even when datasets are anonymized. Membership inference attacks can provide a pathway to deanonymization; knowing that an individual’s data is part of a dataset increases the chances that other techniques or auxiliary information can reveal their identity. Therefore, these two attack types represent the most significant privacy risks associated with the sharing of genomic data and research outcomes. When datasets, even in an anonymized format, are shared, the risk of membership inference inherently overlaps with that of deanonymization. This is especially true when research outcomes, such as detailed statistics about the dataset, are made available, as they can inadvertently expose patterns that allow attackers to identify individual participants.

### Methods for Assessing Privacy Risks in Genomic Data Sharing

Two key methods are used to assess the privacy risks associated with genomic data sharing: the *Likelihood-Ratio Test (LRT)* and the *Hamming Distance* approach [13]. Both methods are commonly employed in membership inference attacks to determine whether an individual’s genome is part of a specific dataset. The Likelihood-Ratio Test is used to evaluate the likelihood that a target individual’s data is part of the case group in a study. The Hamming Distance method, on the other hand, evaluates how closely a target individual’s genome matches any individual’s genome in a dataset.

### Publishing Genomic Research

In genomic research, when published in reputable journals, data availability plays a crucial role in the transparency and reproducibility of scientific findings. The accessibility of different types of data used in these studies varies significantly depending on the nature of the data and the policies of the institutions involved. While genomic research publications make significant efforts to share data with the scientific community, the level of access to different types of data varies depending on privacy concerns, ethical considerations, and institutional policies. The balance between data accessibility and privacy protection is needed to ensure both the advancement of science and the safeguarding of individuals’ sensitive information.

### Data Availability in Genomic Research Publications

Here, we outline the common practices regarding the availability of datasets in genomic research publications:

#### Raw Data Availability

Raw genomic data, which includes the unprocessed sequences and detailed individual-level information, is generally not made publicly available. Researchers or interested parties often need to contact the individual cohorts or the authors directly to request access. This process typically involves submitting an application or an access request, which may require the approval of the authors or the institution responsible for the data. The rationale behind this restricted access is primarily to protect the privacy of individuals whose genomic data is included in the study, as this type of data can be highly sensitive and potentially identifiable.

#### Access to Genome and Biobank Portals

Many genomic studies utilize data from established genome and biobank portals, such as the UK Biobank [14] or the Qatar Biobank (QBB) [15]. The accessibility of data from these sources varies. Some of the data is publicly available, which allows researchers to access it directly without needing special permissions. However, other datasets require a formal request and application process. These applications often need to include a description of the intended research use, and researchers may be required to agree to certain terms and conditions or obtain an accession code. This selective access ensures that the data is used ethically and in accordance with the consent provided by the participants.

#### Availability of Summary Statistics

Summary statistics, which provide aggregated data typically derived from the analysis of the raw genomic data, are more readily available. These statistics might include population-level allele frequencies or associations between genetic variants and specific traits or conditions. They are often made publicly available through repositories like the GWAS Catalog [1], EpiGAD [16], or platforms such as Google Cloud Storage. In some cases, summary statistics are only available after a data transfer agreement is signed, ensuring that the data is shared responsibly.

#### Supplementary Data and Source Files

In addition to the datasets themselves, genomic research publications generally provide supplementary materials, including source data files. These files contain the data used to generate the figures and tables presented in the paper, offering a deeper level of transparency and enabling other researchers to replicate the study’s findings. At least one supplementary data file is typically shared, ensuring that key data underlying the research is accessible to the scientific community.

#### Mention of Data Collection Sites

For studies that involve the collection of new human data, it is common practice for researchers to mention the site of data collection. This information provides context for the data and helps in understanding the population or environment from which the data was derived. Mentioning the data collection site also adds an extra layer of transparency and can be crucial for interpreting the results in the context of population-specific factors.

In summary, while genomic research publications make significant efforts to share data with the scientific community, the level of access to different types of data varies depending on privacy concerns, ethical considerations, and institutional policies. The balance between data accessibility and privacy protection is needed to ensure both the advancement of science and the safeguarding of individuals’ sensitive information.

## Data Sharing Policy Requirements for Genomic Research

In genomic research, ensuring the privacy and security of data is paramount, especially given the sensitive nature of the information involved. The National Institutes of Health (NIH) [2] has established a set of minimum requirements for data sharing to guide researchers in maintaining high standards of data protection and compliance with regulations. These requirements serve as the baseline for responsible data management and sharing practices in the field.

### Informed Consent

Informed consent is the foundation of ethical data sharing in genomic research. Researchers are required to clearly communicate to participants how their data will be used, shared, and protected, including any limitations or conditions for sharing. This process ensures that participants are fully aware of the potential uses of their data and have willingly agreed to these terms.

### De-identification and Anonymization

To protect participant identity, personal identifiers must be removed or obscured from the data. This is achieved through de-identification and anonymization techniques such as data masking or aggregation. These methods prevent direct or indirect identification of participants, thus safeguarding their privacy.

### Data Encryption

Encryption is crucial for protecting data against unauthorized access. This involves implementing strong encryption methods for both stored data and data being transferred. Encryption ensures that data remains confidential and is only accessible to individuals with the necessary decryption keys.

### Access Controls

Access to data is restricted to authorized personnel only. This is managed through the implementation of robust access controls and authentication mechanisms, such as passwords and multi-factor authentication. Role-based access controls ensure that individuals can only access data pertinent to their role within the research project.

### Data Use Agreements (DUAs)

DUAs are formal agreements that specify the terms and conditions under which data can be accessed and used by third parties. These agreements are crucial for ensuring that all parties involved adhere to the agreed-upon data security and privacy practices.

### Audit Trails and Monitoring

Maintaining audit trails is essential for logging access and usage of data. These logs help monitor data access and usage to detect and respond to unauthorized activities promptly. Audit trails contribute to the integrity and accountability of the data management process.

### Compliance with Regulations

Researchers must ensure that their data management practices comply with relevant laws and regulations, such as the Health Insurance Portability and Accountability Act (HIPAA) [17] in the United States and the General Data Protection Regulation (GDPR) [18] in the European Union. Compliance helps protect research participants and institutions from legal and ethical issues.

### Data Sharing Repositories

Data sharing repositories play a critical role in genomic research by providing a platform for data accessibility. These repositories must adhere to FAIR (Findable, Accessible, Interoperable, and Reusable) principles and enforce stringent privacy and security measures. Verification of these repositories to ensure they have appropriate certifications and compliance mechanisms is essential for maintaining trust and integrity in the data sharing process.

By adhering to these NIH guidelines, genomic researchers can ensure that their practices not only advance scientific discovery but also respect and protect the privacy and rights of participants. This balanced approach is crucial for fostering trust and cooperation between researchers and participants, which is essential for the ongoing success of genomic research.

## Methods

In this section, we discuss workflow and survey design for evaluating data sharing and privacy protection in genomic research.

### Proposed Workflow for Privacy Risk Evaluation

To rigorously assess privacy risks associated with data sharing in genomic research, we have developed a structured workflow that follows three key steps as mentioned in the ‘Overview of the Bidirectional Framework’: a questionnaire, a comparative analysis, and feedback to researchers.

The workflow begins with the **Questionnaire** phase, where a targeted survey is conducted to collect detailed information from researchers about their data sharing practices and the privacy protection measures they have implemented. This step is designed to gather insights into the type of data being shared, the level of anonymization, and the risk mitigation strategies in place.

In the **Comparative Analysis** phase, the collected survey data is systematically compared against the National Institutes of Health (NIH) Genomic Data Sharing Policy, which serves as a benchmark for evaluating the adequacy of the privacy protections employed by the researchers. The workflow follows a decision-tree approach (illustrated in Figure 1, where the responses from the survey are analyzed to determine whether they align with NIH guidelines. This step is crucial as it establishes whether the researchers’ practices meet the baseline requirements for ethical data sharing and privacy protection.

**Figure 1:**
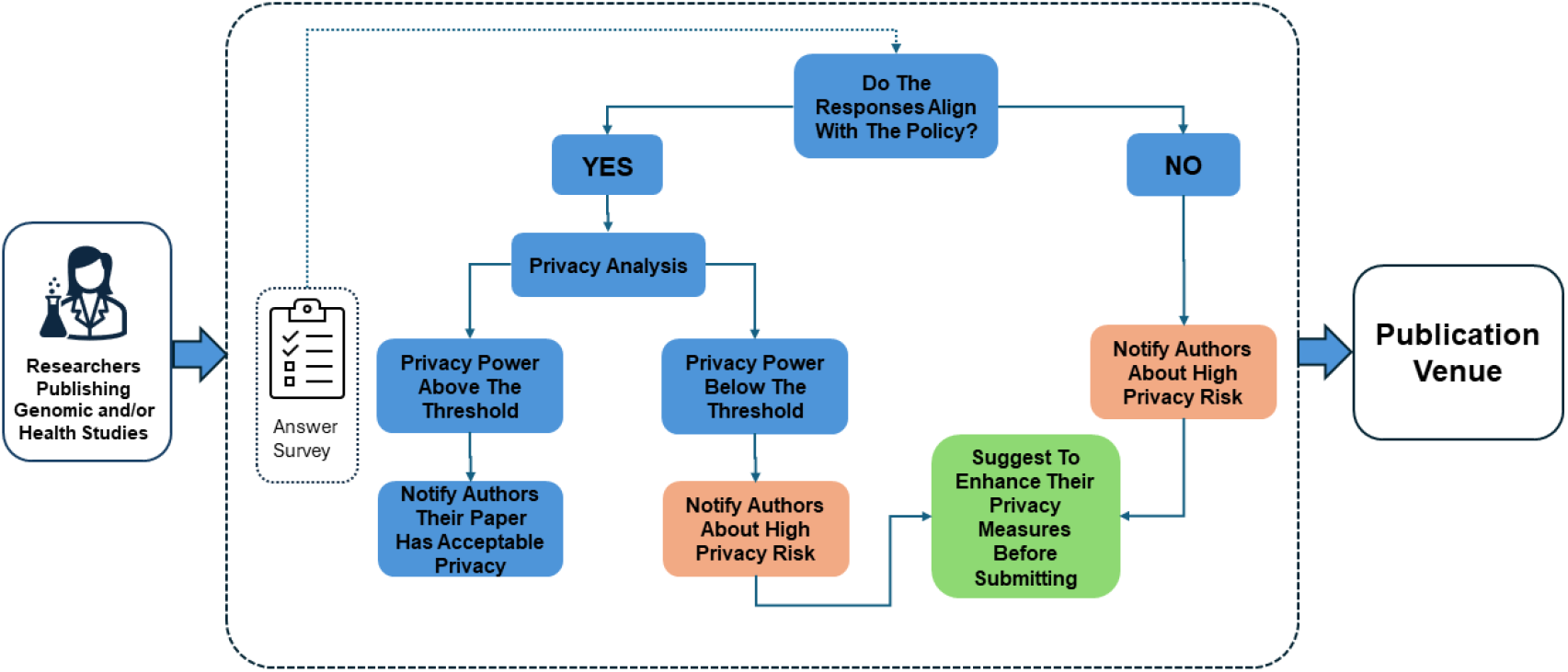
A comprehensive workflow for evaluating privacy risks in genomic data sharing, outlining the three key steps of the proposed bidirectional framework. The process begins with a questionnaire phase where researchers provide detailed information on their data-sharing practices and privacy measures. This is followed by a comparative analysis against the National Institutes of Health (NIH) Genomic Data Sharing Policy to assess compliance and identify potential vulnerabilities. Finally, feedback is provided to researchers, including suggested actions to enhance privacy protection measures before publication. This iterative approach aims to ensure both data accessibility for scientific research and the protection of individual privacy.

If the practices align with the NIH policy, the workflow proceeds to apply a **Membership Inference Attack (MIA) analysis** to assess the risk of re-identification. The power of the MIA is evaluated to determine whether it exceeds a predefined threshold. If the MIA power is below the threshold, indicating a lower risk of re-identification, the study may continue toward submission.

However, if the practices do not align with the NIH data sharing policy, or if they do align but the MIA power is above the threshold, the workflow moves to the **Feedback** phase. At this stage, researchers are notified of the high privacy risk associated with their data sharing practices. They are encouraged to enhance their privacy measures, such as implementing stricter access controls or applying noise addition techniques, before proceeding with submission. If the researchers choose to accept the feedback and strengthen their privacy protections, they are invited to resubmit their work for another round of evaluation.

Alternatively, if researchers opt not to enhance their privacy measures, they may still choose to submit their research, but this is done at their own risk, with full awareness of the potential privacy vulnerabilities. Depending on the actions taken by the researchers, the paper is either submitted with strengthened protections or submitted with the identified risks intact.

This workflow ensures a systematic and iterative approach to evaluating and mitigating privacy risks, promoting the responsible sharing of genomic data in a manner that balances the need for scientific transparency with the imperative to protect individual privacy.

### Survey Design for Data Sharing and Privacy Protection

A well-structured survey is a vital component of the proposed workflow, serving as the primary tool for collecting detailed information on the data sharing practices and privacy protection measures implemented by researchers in genomic studies. The survey is designed to gather targeted insights into the types of data shared, the specific strategies used to mitigate privacy risks, and the level of adherence to the National Institutes of Health (NIH) guidelines. By asking a series of focused questions, the survey aims to uncover both current practices and potential gaps in privacy protection.

The survey, consisting of ten questions,^*^ begins by determining the type of data involved, such as human genome data or clinical data, to assess the sensitivity of the information being shared. It then probes the data sharing practices used during the research process to identify any inherent privacy risks. For instances where data sharing occurred, the survey asks about the risk mitigation strategies employed, such as de-identification or restricted access.

In addition, the survey examines the privacy protection measures used, including techniques like encryption and anonymization, and seeks detailed explanations of how these measures were implemented to assess their robustness. It evaluates whether summary statistics and/or data from the research were made publicly available and to what extent, highlighting the potential for indirect re-identification risks.

To further assess the practical application of privacy measures, the survey also asks researchers to provide a sample of the data or statistics they are willing to share along with their paper for publication. By requesting a sample of the data or statistics intended for sharing, the survey not only gathers comprehensive data on current practices but also enables a practical examination of potential privacy risks.

Finally, the survey explores whether there are plans to enhance privacy measures in future data sharing efforts, emphasizing a proactive and forward-looking approach to data protection.

The insights gained from the survey directly inform the workflow, guiding the evaluation process to ensure that genomic research adheres to the highest standards of privacy protection while balancing the need for open scientific inquiry.

## Results

In this section, we evaluate the privacy risks associated with genomic research using two case studies where researchers share additional materials along with their publications. We consider two different scenarios: in the first scenario, researchers share only the summary statistics of their research; in the second scenario, they share both raw genomic data and the associated statistical results. Below, we delve into each scenario to assess the potential privacy risks.

### Case study 1: Privacy Risk Evaluation in Genomic Research that shared GWAS Statistics

In this case study, we examine a genomic research article published in *Nature*^†^ that used human genome data from various biobanks to investigate genetic factors and cellular mechanisms. This case study illustrates the use of a structured workflow to evaluate the privacy risks associated with the sharing genomic research finding and statistics, particularly in the context of data sharing policies and re-identification risks.

#### Data Sharing and Privacy Protection Measures

The researchers in this study followed rigorous data-sharing practices consistent with the regulations of the biobanks from which the data was sourced. Data was shared through various controlled-access portals, ensuring that access was restricted and provided only to research projects that had obtained the necessary ethical approvals. This practice aligns with the National Institutes of Health (NIH) baseline [2] requirements for responsible data sharing.

The researchers implemented multiple risk mitigation strategies to protect participant privacy. These included the de-identification of individual-level health, lifestyle, demographic, and genetic data. Anonymization techniques were applied to remove or obscure personal identifiers, and stringent data access protocols were enforced. These protocols required approval from ethics review committees and data access committees before any data could be accessed or shared.

#### Availability of Research Statistics

The study’s summary statistics were partially made available to the public. Overall and population-level Genome-Wide Association Studies (GWAS) summary statistics were shared on the GWAS Catalog, a public repository for genomic data. Other data, however, were accessible only upon reasonable request or through specific data access systems, ensuring that sensitive information remained protected.

#### Evaluation Against NIH Policy

The study’s data sharing practices were evaluated against the NIH data sharing policy using proposed workflow. The survey responses indicated that the practices aligned with NIH guidelines, ensuring that informed consent, de-identification, encryption, and access controls were appropriately implemented.

However, during the Membership Inference Attack (MIA) analysis using the Likelihood-Ratio Test, the MIA power was found to be high, exceeding 50% when more than 250 SNPs were shared in the case group (as shown in Figure 2). For this analysis, we specifically considered p-values less than 0.05 in the case group to identify underrepresented groups, as these lower p-values indicate significant associations with rare traits or conditions. The Likelihood-Ratio Test was then applied to assess the potential for re-identification based on these significant associations.

**Figure 2:**
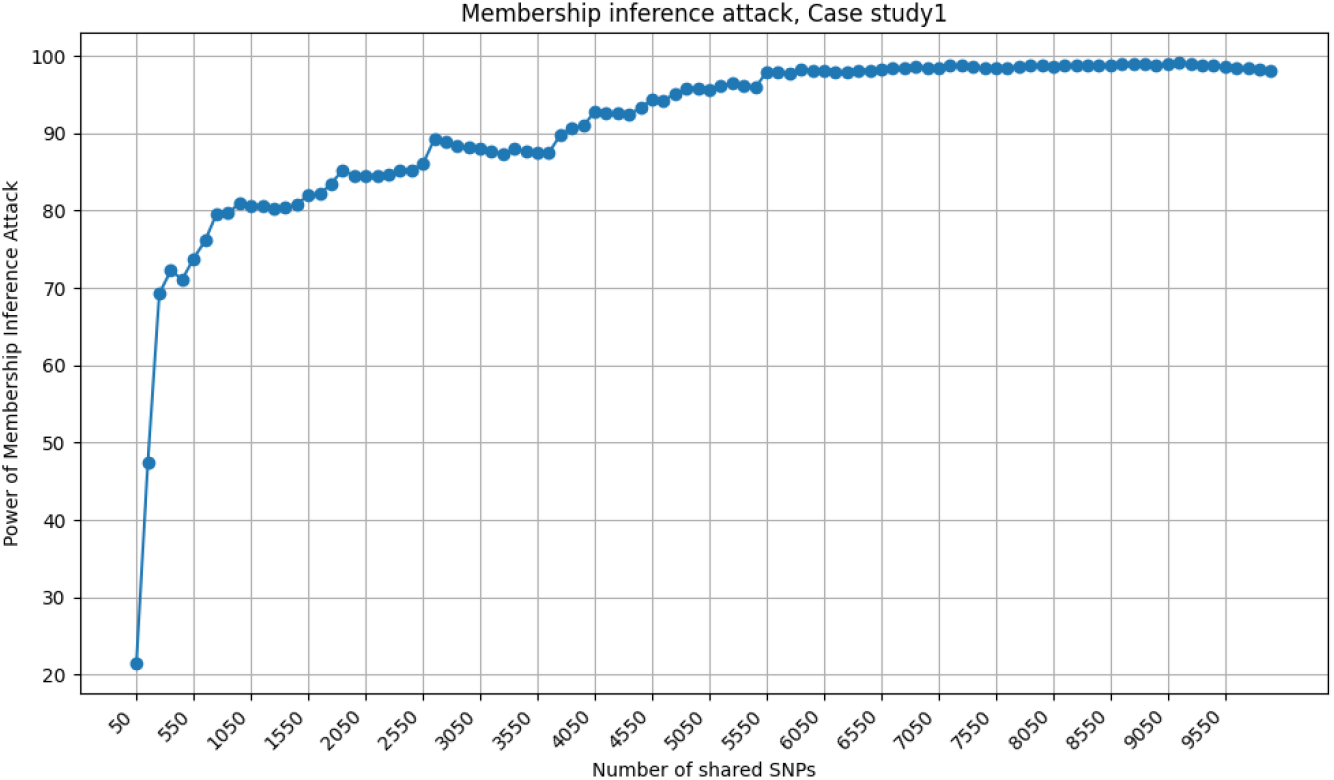
Analysis of the power of a Membership Inference Attack (MIA) in Case Study 1, which involves the sharing of GWAS summary statistics. The plot illustrates the increased risk of re-identification associated with the sharing of more than 250 Single Nucleotide Polymorphisms (SNPs) in a given case group. As shown, the power of the attack exceeds 50%, indicating a significant privacy threat, particularly when large datasets or datasets involving rare genetic traits are shared. The results underscore the need for stringent privacy protection measures to mitigate re-identification risks.

#### Recommendations for Privacy Enhancement

Given the high MIA power observed in this case, the authors will be advised to consider additional privacy protection strategies. These could include applying noise addition techniques. If the authors chose to accept these suggestions, they could then resubmit their data for another round of privacy risk evaluation. Otherwise, they were informed that submitting the research in its current form could involve significant privacy risks, potentially compromising the confidentiality of the study participants.

### Case Study 2: Privacy Risk Evaluation in a Genomic Research that shared Both Raw Data and Statistics

In the second case study, we explore a scenario where researchers have shared both raw genomic data and accompanying statistical results as part of their publication. This practice, while enhancing transparency and fostering collaboration, introduces significant privacy risks. Even when researchers strictly adhere to established data sharing policies, such as those outlined by the NIH, the potential for re-identification of individuals remains a concern, particularly through membership inference attacks (MIA).

Membership inference attacks involve determining whether an individual’s data is included in a particular dataset. In this scenario, we assume that the researchers’ data sharing practices are fully aligned with NIH guidelines, meaning that they have implemented all recommended safeguards, such as informed consent, de-identification, and data access controls.

To analyze the risk of MIA, we used a publicly available dataset from the OpenSNP [19], which includes phenotype information about the individuals. From this dataset, we created two groups: a case group consisting of individuals with a specific genetic trait or condition and a control group without that trait.

To assess the risk of MIA in this scenario, we applied a membership inference attack using a Hamming distance approach, as explained in the previous sections.

Figure 3 illustrates the power of a membership inference attack when applied to this shared genomic dataset. The plot reveals that as the number of shared SNPs increases, the power of the attack approaches its maximum value. Even after the initial sharing of around 10 SNPs, the power of the attack rises sharply, reaching 100% by sharing more than 160 SNPs. This indicates a high likelihood that an attacker could successfully infer the membership of individuals in the dataset.

**Figure 3:**
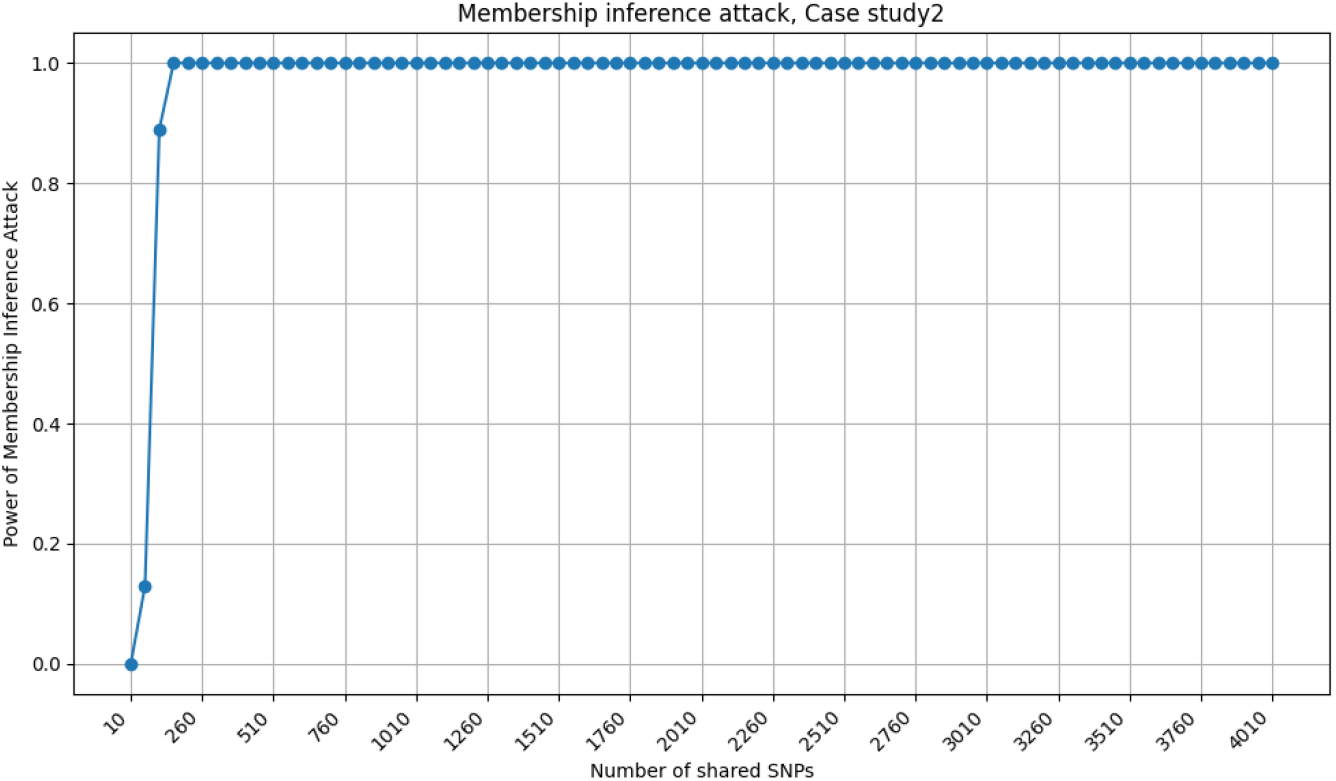
Detailed evaluation of the power of a Membership Inference Attack (MIA) in Case Study 2, where both raw genomic data and associated statistical results are shared. This plot demonstrates the relationship between the number of SNPs shared and the likelihood of successful re-identification by an attacker. The attack power sharply increases after the sharing of more than 50 SNPs and reaches its maximum when 250 or more SNPs are shared. These findings highlight the substantial privacy risks inherent in sharing raw genomic data, even when institutional guidelines are followed, suggesting the need for advanced privacy-preserving techniques.

The findings underscore the substantial privacy risks associated with sharing both raw genomic data and statistical information. Even with compliance to NIH guidelines, the risk of re-identification remains high, suggesting the need for additional safeguards and more advanced privacy-preserving techniques to protect participant confidentiality.

## Conclusion

This study presents a robust framework designed to support authors of genomic research papers in navigating the growing concerns surrounding privacy in genomic research, particularly in the context of data sharing. By integrating a detailed workflow with a questionnaire and privacy risk assessments, our framework provides a practical approach to ensuring that the benefits of data sharing are realized without compromising the privacy of individuals. The framework’s emphasis on comparing researchers’ practices with NIH guidelines and conducting Membership Inference Attack (MIA) analyses provides a comprehensive method for identifying and mitigating potential re-identification risks.

The case studies included in this paper illustrate the complexities and challenges that authors face in balancing the need for data accessibility with the responsibility to protect participant privacy. These examples demonstrate the real-world applicability of our framework, showing how it can be used to guide authors in making informed decisions about their data sharing practices, ultimately leading to more secure and privacy-conscious research outputs.

In conclusion, the framework developed in this study provides a valuable tool for researchers and authors, helping them navigate the complex landscape of genomic data sharing. It offers clear guidance on maintaining the delicate balance between advancing scientific discovery and protecting individual privacy, ensuring that the trust placed in genomic research by participants and the public is both earned and preserved. By providing this support, we aim to empower authors to share their research findings responsibly and ethically, fostering greater collaboration and progress in the field.

## Acknowledgments

The work was partly supported by the National Library of Medicine of the National Institutes of Health under Award Number R01LM013429, the National Science Foundation (NSF) under grant numbers 2141622, 2050410, 2200255, and OAC-2112606, and Cisco Research.

## Footnotes

* Due to space constraints, we did not provide a detailed description of each question.

† For the purpose of privacy, we intentionally do not cite the specific research paper.

